# Defining data-driven primary transcript annotations with *primaryTranscriptAnnotation* in R

**DOI:** 10.1101/779587

**Authors:** Warren D. Anderson, Fabiana M. Duarte, Mete Civelek, Michael J. Guertin

**Affiliations:** Center for Public Health Genomics, University of Virginia, Charlottesville, Virginia, United States of America; Department of Stem Cell and Regenerative Biology, Harvard University, Cambridge, Massachusetts, United States of America; Department of Biomedical Engineering, University of Virginia, Charlottesville, Virginia, United States of America; Biochemistry and Molecular Genetics Department, University of Virginia, Charlottesville, Virginia, United States of America

**Author notes:** co-senior authors (Mete Civelek, Michael J. Guertin).

**Keywords:** PRO-seq/GRO-seq analysis, transcript annotation, RNA Polymerase Pausing

## Abstract

Nascent transcript measurements derived from run-on sequencing experiments are critical for the investigation of transcriptional mechanisms and regulatory networks. However, conventional gene annotations specify the boundaries of mRNAs, which significantly differ from the boundaries of primary transcripts. Moreover, transcript isoforms with distinct transcription start and end coordinates can vary between cell types. Therefore, new primary transcript annotations are needed to accurately interpret run-on data. We developed the `primaryTranscriptAnnotation` R package to infer the transcriptional start and termination sites of annotated genes from genomic run-on data. We then used these inferred co-ordinates to annotate transcriptional units identified *de novo*. Hence, this package provides the novel utility to integrate data-driven primary transcript annotations with transcriptional unit coordinates identified in an unbiased manner. Our analyses demonstrated that this new methodology increases the sensitivity for detecting differentially expressed transcripts and provides more accurate quantification of RNA polymerase pause indices, consistent with the importance of using accurate primary transcript coordinates for interpreting genomic nascent transcription data.

**Availability:** **https://github.com/WarrenDavidAnderson/genomicsRpackage/tree/master/primaryTranscriptAnnotation**

## Introduction

Quantification of nascent transcription is critical for resolving temporal patterns of gene regulation and defining gene regulatory networks. Processed mRNA levels are influenced by numerous factors that coordinate the mRNA production and degradation rates (Blumberg et al., 2019; Honkela et al., 2015). In contrast, the levels of nascent RNAs reflect the genome-wide distribution of transcriptionally engaged RNA polymerases at the time of measurement (Wissink et al., 2019). Precision run-on sequencing (PRO-seq, Kwak et al. (2013)) and global run-on sequencing (GRO-seq, Core et al. (2008)) have standardized experimental protocols and are commonly used to quantify nascent transcription (Lopes et al., 2017; Mahat et al., 2016). Genome-wide analyses of nascent transcription require accurate annotations of gene boundaries. While ongoing efforts aim to increase the quality of genome annotations (Haft et al., 2018), existing gene annotations are inadequate for both quantifying nascent transcripts and determining the RNA polymerase location relative to gene features. Analyses of run-on data indicates that annotated transcription start sites (TSSs) are often inaccurate (Link et al., 2018). Similarly, it is well established that transcription extends beyond the 3′ polyadenylation region (Proudfoot, 2016), thereby rendering transcription termination sites (TTSs) distinct from annotated mRNA ends. Identifying more accurate TSSs and TTSs for primary transcripts is important for accurate transcript quantification from run-on data. Experimental techniques such as 5′ GRO-seq, PRO-cap, and Start-seq can directly estimate TSS coordinates (Link et al., 2018; Mahat et al., 2016; Scruggs et al., 2015), however, data-driven methods for improved annotations are of considerable practical interest.

Efforts in *de novo* transcript identification from run-on data have partially addressed problems related to TSS/TTS annotation. The R package `groHMM` and the command line tool `HOMER` identify transcriptional unit (TU) coordinates and have been successfully applied to run-on data (Chae et al., 2015; Heinz et al., 2010). However, these existing methods do not facilitate the assignment of gene identifiers to the identified TUs, which are generically defined by their chromosomal coordinates.

Here we present the R package `primaryTranscriptAnnotation` for annotating primary transcripts. We directly infer TSSs and TTSs for annotated genes, then we integrate the identified coordinates with TUs identified *de novo*. Our improved annotations increase the sensitivity and accuracy of detecting differential transcript expression and quantifying RNA polymerase pausing. This package improves precision in analyses of critical phenomena related to transcriptional regulation and can be easily incorporated into standard genomic run-on analysis workflows.

## Description

We distinguish two related tasks performed by our package: (1) Integration of run-on data and existing gene annotations to refine estimates of TSSs and TTSs, and (2) combining the results of the first task with the results of an unsupervised TU identification method (`groHMM` or `HOMER`) to annotate the TUs. We accept the data-driven annotations from (1) as a ‘ground truth’ and we use these coordinates to segment and assign identifiers to the *de novo* TUs (Figure 1a). For demonstration of the package functions, we use PRO-seq data from adipogenesis time-series experiments available through GEO accession record GSE133147. Extensive implementation details are provided in the vignette associated with the publicly available R package.

**Fig. 1.**
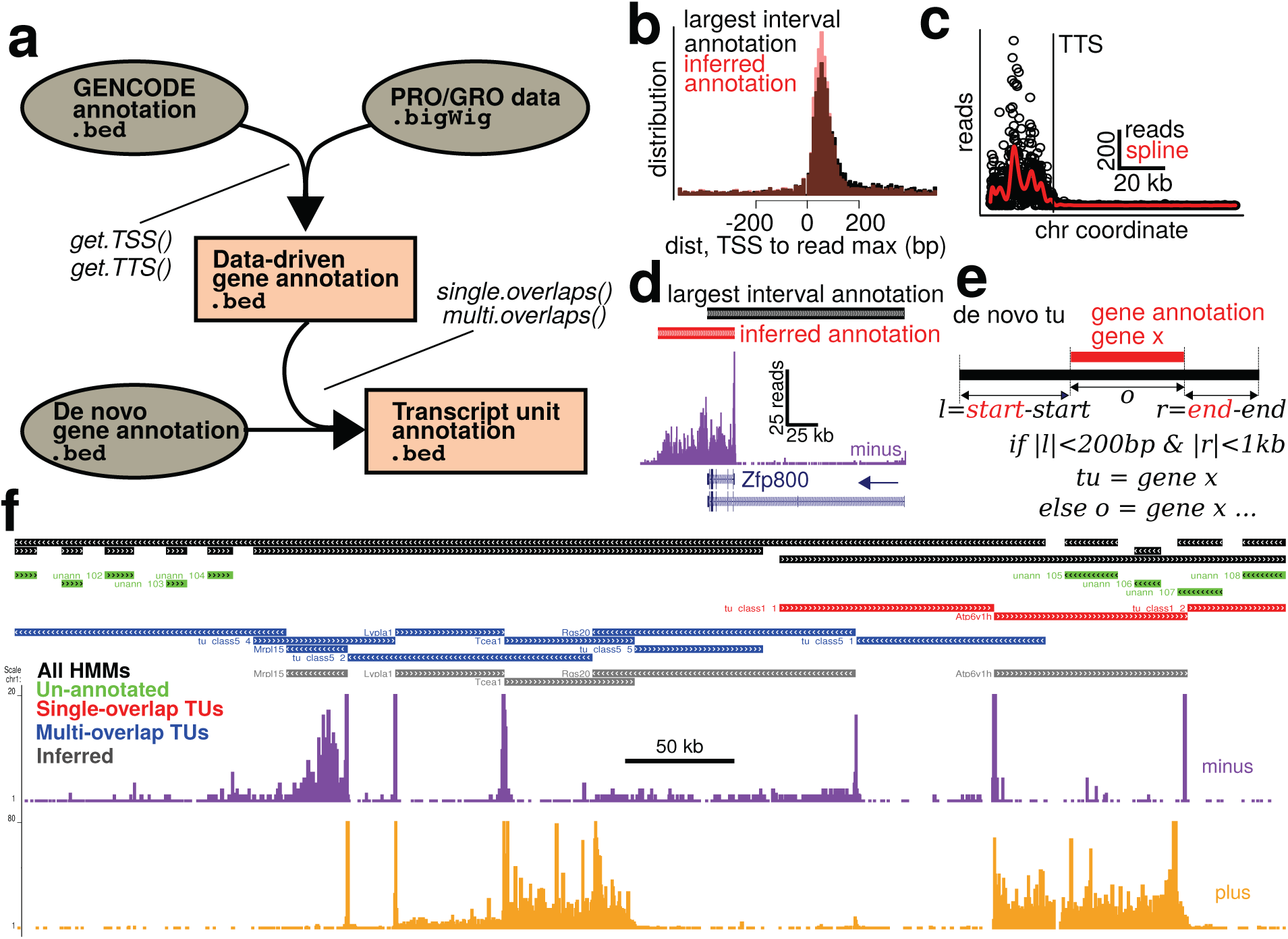
The primaryTranscriptAnnotation package accurately annotates gene features and assigns gene names to transcriptional units. (a) Aligned run on data and gene annotations are inputs to redefine gene annotations. *De novo*-identified transcription units can be assigned gene identifiers using refined gene annotations. (b) Promoter-proximal paused RNA polymerases are more constrained to the canonical 20-80 bases downstream of the refined transcription start sites, compared to conventional annotations. (c) TTS inference involves 1) detection of higher density peaks in the 3′ end of the gene, corresponding to the slowing of RNA polymerase and 2) determining the genomic position when the read density decays towards zero. (d) These methods generate improved TSS and TTS estimates for the Zfp800 gene. (e) Annotation of *de novo*-defined transcription units with gene identifiers is based upon degree of overlap. (f) This approach produces gene boundaries with improved accuracy, while maintaining gene identifiers.

### Data-driven gene annotation

We used GENCODE gene annotations as a reference point for inferring TSSs and TTSs (Harrow et al., 2012). To infer TSSs, we considered all first exons of each gene isoform and defined the TSS as the 5′ end of the annotated first exon that contains the maximal read density within a specified range downstream. Such regions of peak read density, typically between 20 and 80 bp downstream from a TSS, exist at RNA polymerase ‘pause sites’ (Kwak et al., 2013). To evaluate the performance of our TSS identification method, we compared our inferences to the GENCODE annotations. We considered the ‘largest interval’ annotation for each gene by taking the most upstream start coordinates and most downstream end coordinates from the GENCODE annotation file. Figure 1b and Figure S1 show the distance between the gene start coordinate and the nearest read peak within 500 bases. Consistent with paused RNA polymerase accumulation in close proximity and downstream of transcription initiation sites, the distribution of highest RNA polymerase densities is more focused immediately downstream for the refined TSS annotation as compared to the ‘largest interval’ annotation (Figure 1b and Figure S1).

To infer TTSs, we examined evidence of transcriptional termination in regions extending from a 3′ interval of the gene to a selected number of base pairs downstream of the most distal annotated gene end (Figure S2). We based our method for TTS inference on data demonstrating that transcription rates are attenuated at gene ends (Lian et al., 2008). This phenomena is manifested as elevated polymerase density at the gene end relative to the middle of the gene body (Fong et al. (2015), see also Figure S3). We considered the possibility that the identification of TTSs could be influenced by distal enhancers or downstream promoters that are defined by bidirectional transcription (Core et al., 2014). To reduce the likelihood of such occurrences, we used an exponential model that incorporates the distance to the downstream gene in order to define a search region for identifying read density peaks corresponding to elevated polymerase density at gene ends (Figure S2). We defined the TTSs by binning the search regions, counting reads within the bins, fitting smooth spline curves to the binned counts, identifying peaks in the curves, and detecting points at which the curves decay from the peak towards zero (Figure 1c, Figure S3). The results of our TTS identification procedures show that data-driven annotations result in considerably larger gene coordinates as compared to conventional mRNA annotations, which is consistent with RNA polymerases transcribing beyond the polyadenylation and cleavage site (Figure 1d, Figure S4d,e).

### Annotation of *de novo* transcriptional units

While data-driven gene coordinates provide an improvement over conventional annotations, it can be advantageous to analyze run-on data in the context of TUs identified in an unbiased manner (Chae et al., 2015). Given both *de novo* TUs and a trusted gene annotation, `primaryTranscriptAnnotation` combines these information sources to annotate the TUs so that TUs overlapping genes are assigned conventional gene names (Figure 1e). We separately considered either TUs that overlapped single genes or TUs that overlapped multiple genes (Figures S5,S6). If the start and end of a TU were within user-defined distances to the TSS and TTS of an overlapping gene, the TU was assigned the identifier for the overlapping gene. If the TU did not extend to the gene boundaries, the TU was extended to match the trusted gene annotation. TUs that did not overlap any genes were marked as unannotated. Figure 1f shows PRO-seq reads along with `groHMM` annotations (black), TSS/TTS inference annotations (grey), and annotations based on combining the results of `groHMM` and TSS/TTS inference (green, red, blue). Combining both the data-driven gene annotation and transcript unit annotation methods provides more accurate transcript boundaries and retains gene identifier information that can be used in down-stream applications.

### Improved sensitivity for detecting gene expression changes and RNA polymerase pausing

We annotated PRO-seq data with both ‘largest interval’ gene coordinates and inferred primary transcript coordinates (e.g., see Figure 1d). Principal component analysis shows that TU/gene annotations based on our novel approach preserve the data structure observed using conventional largest interval annotations (Figure S7). However, when we examined differential expression by applying a likelihood ratio test to our adipogenesis time-series data, the results showed that using the inferred gene coordinates resulted in more sensitive detection of differential expression (Supplemental Results and Figure S8). This finding demonstrates that our inferred annotations result in enhanced sensitivity for detecting differential expression of run-on data.

Promoter proximal polymerase pausing is a pervasive phenomena in eukaryotic gene regulation, and has been implicated in numerous biological functions including development, environmental response, and cell differentiation (Adelman and Lis, 2012; Duarte et al., 2016; Scheidegger and Nechaev, 2016). To determine if our inferred annotations confer an improvement for detecting RNA polymerase pausing, we computed ‘pause indices’ based on data from inferred coordinates and largest interval annotations. The pause index is a ratio of read density near the TSS to read density within the gene body (Min et al., 2011). We defined the pause region as 20-80 bp downstream of the TSS and the gene body region as the interval from 500 bp downstream of the TSS to the gene end. Our analyses revealed that applying inferred gene coordinates results in significantly larger pause indices (Figure S9). This suggests that our methods enhance the sensitivity for detecting and quantifying pausing at a genome-wide scale.

## Discussion

We describe `primaryTranscriptAnnotation` and illustrate its utility with analysis examples. The package is publicly available, requires minimal dependencies, and is easy to use. The package also includes the data necessary to reproduce all analyses presented in this paper. The associated vignette and Supplementary Materials contains code for illustrating the package functionality and reproducing all analyses. Because run-on data is critical to defining transcriptional regulatory mechanisms, and because existing gene annotations are suboptimal for mapping run-on data, `primaryTranscriptAnnotation` will be generally useful for investigations into the mechanisms of transcription.

## Data availability

Raw sequencing files and processed *bigWig* files are available from GEO accession record GSE133147.

## ACKNOWLEDGEMENTS

Thanks to Dr. Thurl Harris for providing the 3T3-L1 cells. Thanks to the laboratory of Dr. Chongzhi Zang, as well as the Civelek and Guertin laboratories, for advice and discussion. Thanks to Dr. Yongde Bao with the UVA Genome Analysis and Technology Core for sequencing our PRO-seq libraries.

## Supplemental

### Supplemental Materials

#### Cell culture and adipogenesis

We performed adipogenesis experiments with 3T3-L1 cells grown in high glucose (4500 mg/L) Dulbecco’s Modified Eagle Medium (DMEM), 10% newborn calf serum, 1% fetal calf serum, 100 U/mL penicillin G sodium, and 100 *µ*g/mL streptomycin sulfate. During expansion of the fibroblast adipocyte precursors, we passaged the cells prior to confluence. For the final passage before differentiation, we allowed the cells to reach confluence. We differentiated the adipocyte precursors into mature adipocytes by applying a standard differentiation media to the cells three days following confluence. The differentiation media included high glucose DMEM, 10% fetal calf serum, 100 U/mL penicillin G sodium, 100 *µ*g/mL streptomycin sulfate, 0.25 U/mL insulin, 0.5 mM 3-isobutyl-1-methylxanthine (IBMX), and 0.25 *µ*M Dexamethasone (Adamson et al., 2015). Differentiation proceeded for up to four hours for the experiments reported in this manuscript. We sampled cells undergoing adipogenesis at the following time points with respect to the application of differentiation media: time 0 (no differentiation media), 20 min, 40 min, 60 min, 2 hr, 3 hr, and 4 hr. We obtained three replicates at each of the seven sample time points.

#### PRO-seq experiments

We performed PRO-seq experiments as described previously (Sathyan et al., 2019). In brief, we washed the cells with an ice-cold wash buffer (10 mM Tris-HCl pH 7.5, 10 mM KCl, 150 mM Sucrose, 5 mM MgCl_2_, 0.5 mM CaCl_2_, 0.5 mM DTT, 0.004 units/ml SUPERaseIN RNase inhibitor (Invitrogen), and Protease inhibitors (complete, Roche)). We then permeabilized the cells for 3 min with a permeabilization buffer (10 mM Tris-HCl pH 7.5, 10 mM KCl, 250 mM Sucrose, 5 mM MgCl_2_, 1 mM EGTA, 0.05% Tween-20, 0.1% NP40, 0.5 mM DTT, 0.004 units/ml SUPERaseIN RNase inhibitor, and Protease inhibitors) and washed again with wash buffer. Next, we suspended the cells in a freezing buffer (50 mM Tris-HCl pH 8, 5 mM MgCl_2_, 0.1 mM EDTA, 50% Glycerol and 0.5mM DTT) and isolated ∼ 3 − 5 × 10^6^ cells per 50 *µ*L of glycerol buffer for run-on experiments, rapidly froze the suspension in liquid Nitrogen, and stored the cells at -80°C for ∼1 week. We prepared the PRO-seq libraries as described previously (Sathyan et al., 2019). We included random 8 bp unique molecular identifiers (UMIs) at the the 5′end of the RNA adapters that are ligated to the 3′ ends of the nascent transcripts to allow for removal of PCR duplicates (Fu et al., 2018; Kivioja et al., 2012). We sequenced pooled libraries on an Illumina NextSeq500 with 75 bp single end reads.

#### PRO-seq analysis

Our scripts for basic alignments and file conversions are publicly available: https://github.com/WarrenDavidAnderson/genomeAnalysis/tree/master/PROseq. We clipped the sequencing adapters using `cutadapt` (Martin, 2011), removed UMI duplicates, trimmed the first 8 UMI bases using `fastx_trimmer` (Gordon, 2010), and obtained the reverse complement of each read using `fastx_reverse_complement` (Gordon, 2010). We aligned the processed reads to the *mm10* mouse genome build using `bowtie2` and filtered reads with a low probability of unique alignment using `samtools` (-q 10) (Langmead et al., 2009; Li et al., 2009). We strand-separated the aligned bam files using `samtools`. We used `seqOutBias` (Martins et al., 2018) to simultaneously shift the alignments to represent the 3′end of the RNA and convert the BAM to the bigWig format. We merged the strand-specific bigWig files from all replicates using the UCSC Genome Browser Utilities (Kent et al., 2010). We also generated bigWig files corresponding to individual replicates for analysis of the time series data. We used the `bigWig` R package to load and operate upon bigWig files (Martins, 2015). We used `Bedtools` for overlapping transcript coordinates, defined by inferred transcription start sites (TSSs) and transcription termination sites (TTSs), with transcript coordinates defined by conventional annotations (Quinlan and Hall, 2010).

#### De novo transcript identification

We used the `groHMM` R package to identify transcriptional units in an unbiased manner from read data merged across time points (Chae et al., 2015). Because `groHMM` supports parameter tuning for performance optimization, we varied key analysis parameters and evaluated the results. We varied the log probability of a transition from the transcribed to the untranscribed state and the read count variance in the untranscribed state (LtProbB and UTS, respectively). For this analysis, we used the inferred primary transcript annotations, generated using `primaryTranscriptAnnotation` to determine TSSs and TTSs, to evaluate the performance of each set of transcription units (TUs) identified from a given Hidden Markov Model (HMM) parameterization. The evaluateHMMInAnnotations() function uses the inferred gene annotations to document ‘merge errors’ and ‘dissociation errors’. Merge errors occur when a given TU overlaps multiple gene annotations. Dissociation errors occur when multiple TUs overlap a given gene annotation. Distinct groHMM parameterizations produced varying degrees of merge and dissociation error. In addition to evaluating merge and dissociation errors, we evaluated HMM sensitivity by looking at how many reads mapped to regions of the genome that were not identified as transcribed. We selected the HMM with the lowest read count outside of HMM-defined transcribed regions. This HMM had relatively low merge error and dissociation error (both within the lowest error quartile). The code for these analyses is publicly available: https://github.com/WarrenDavidAnderson/genomeAnalysis/tree/master/groHMMcode.

#### Principal component analysis and differential expression analysis

We used `DESeq2` to identify ‘size factors’ to normalize the individual PRO-seq sample read data based on sequencing depth (Love et al., 2014). We then applied a variance stabilizing logarithmic transformation using rlogTransformation(). We applied principal component analysis (PCA) using the singular value decomposition-based R function prcomp(). For differential expression analysis, we identified genes with statistically significant temporally varying expression levels using a likelihood ratio test with the `DESeq2` function DESeq(). This test compares the likelihood of a model incorporating time to the likelihood of a null model in which time is not considered.

#### Pause region analysis

We quantified promoter-proximal polymerase pausing by defining a pause index as the ratio of the PRO-seq read signal in the vicinity of the TSS to that signal within the gene body (Min et al., 2011). We defined the pause region between 20 bp and 80 bp downstream of the TSS. We defined the gene body region from 500 bp downstream of the TSS to the gene end. Within each region, we computed read densities and we took the pause/body density ratio as the pause index.

#### Code availability

Code for all analyses associated with this manuscript can be found here: https://github.com/WarrenDavidAnderson/manuscriptCode/tree/master/primaryTranscriptAnnotation_code.

### Package details

Here we describe methodological details pertaining to our implementation of analyses performed by the `primaryTranscriptAnnotation` R package. This supplemental section is focused on methodological details, whereas implementation details are described extensively in the package vignette https://github.com/WarrenDavidAnderson/genomicsRpackage/blob/master/primaryTranscriptAnnotation/primaryTranscriptAnnotation-vignette.pdf.

#### Data-driven primary transcript annotation

The first problem addressed by the `primaryTranscriptAnnotation` R package is the data-driven inference of primary transcript coordinates. This problem is addressed by separately identifying TSSs and TTSs with the use of conventional annotations for constraining the regions within which to examine the presence of TTSs/TTSs.

#### Filtering for unexpressed transcripts

To identify transcripts with arbitrarily low expression, we used the read.count.transcript() function. This function determines the transcript with the highest read count for each gene and returns that read count along with the associated read density. Based on analyses of transcript read count and read density distributions, the user can select thresholds below which expression appears to be negligible.

#### Data-driven inference of TSSs

To empirically identify TSSs, we assumed that TSSs are proximal to sites at which polymerase pausing is observed. This assumption is based on a wealth of genome-scale data documenting promoter-proximal pausing (Adelman and Lis, 2012). These data indicate that pausing occurs 20-80 bp downstream of transcriptional initiation. We also assumed that the TSS for each gene is at an annotated first exon for that gene. Accounting for the strand specificity of gene annotations, we defined the TSS as the 5′ end of the exon 1 isoform with the highest density in the region between 20 bp and 120 bp downstream. This analysis is implemented by get.TSS().

#### Filtering for overlapping genes

Annotated genes occasionally overlap with the coordinates defined for other annotated genes. For instance, the end of an upstream gene can be annotated to a coordinate beyond the start of a downstream gene. This can result in confounding gene expression quantification by erroneously counting the same reads for multiple genes. Our package includes a function for identifying such overlaps (gene.overlaps()). The user can then manually evaluate the gene annotations using a visualization tool such as a genome browser (Kent et al., 2002).

#### Data-driven inference of TTSs

To empirically identify TTSs, we assumed that transcription termination occurs within a region including the most down-stream annotated end of all gene isoforms and extending kilobases beyond. We addressed two tasks related to TTS identification: (1) defining the region within which to search for transcriptional termination, and (2) identifying termination within this region.

To define the search region for TTS evaluation, we started by selecting a percentage of the gene end (e.g., 20%). We defined the start of the search region as the difference between the annotated gene end and the gene length multiplied by the specified percentage (see Figure S2c, term *d*_*e*_):

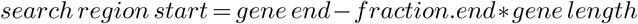

The next step is to specify an upper limit on the total distance from the annotated gene end that could be considered for the TTS search region. This is user specified number of bases (Figure S2c, term *d*_*t*_). We examined whether there were any other gene TSSs in this region between the search region start and the user defined limit. If there was a TSS within the *d*_*t*_ region, we documented the distance between the limit and the TSS. We restricted the search space for TTS evaluation by this value, thus the term *d*_*c*_ is referred to as the clip distance (Figure S2c). Note that the clip distance is dependent on the user defined upper limit.

Experimental data showed that attenuated rates of transcription at gene ends are associated with elevated levels of polymerase density (Lian et al., 2008). Based on this finding, we identified peaks of polymerase density at gene ends for determining where transcription termination is likely to occur. We defined the peak search region *d*_*s*_ as a sub-region of *d*_*a*_ = *d*_*t*_ − *d*_*c*_ (Figure S2c). This is because the identification of gene end polymerase peaks could be affected by distal enhancers that exhibit bidirectional transcription within very large search regions. We assumed that the peak search region should be a large fraction of *d*_*a*_ for genes with the greatest numbers of clipped bases, because such cases occur when the conventional gene ends are proximal to downstream identified TSSs. Thus, in such cases, the entire *d*_*a*_ region should be included for analysis of the gene end peak. The same logic applies for genes with substantially fewer clipped bases, and correspondingly larger *d*_*a*_ regions; in such cases the peak search regions should be smaller proportions of *d*_*a*_. Hence, we defined the gene end peak search region *d*_*s*_ as a function of the clip distance, *d*_*c*_: *d*_*s*_ = *d*_*a*_ *\* f* (*d*_*c*_). The term *f* (*d*_*c*_) gives a value for weighting *d*_*a*_ that is dependent on the clip distance:

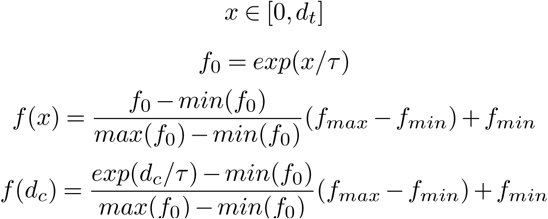

where *τ* is a distance constant defining the rate of exponential decay, *f*_*max*_ is the maximum weight, and *f*_*min*_ is the minimum weight. The terms *f*_*max*_ and *f*_*min*_ should be between zero and one so that the outcome is a fraction. Figure S2b shows the form of the function with dashed lines corresponding to *d*_*c*_ = *τ* = 50 kb, *f*_*min*_ = 0.3, and *f*_*max*_ = 1. This method is used to define the gene end peak search region, *d*_*s*_.

Given the coordinates of the TTS search region and the peak search region, the next task is to identify the TTS within this region. We operationally defined the TTSs by binning the search regions, counting reads within the bins, fitting smooth spline curves to the binned counts, identifying gene end peaks of smooth curves, and detecting points at which the curves decay from the gene end peaks towards zero (Figure S3a,b). For this analysis, we applied constraints on the number of bins used for quantification, and the number of knots for the spline fits, as described in the vignette. For the spline fit within the peak search region, we identified the largest peak and then we determined the point at which the trace decays to a point below a specified threshold of the peak height. The function get.TTS() implements the TTS evaluation procedures described above.

#### Annotation of *de novo* transcriptional units

The second problem addressed by the `primaryTranscriptAnnotation` R package is the annotation of transcriptional units (TUs), identified *de novo*, using primary transcript coordinates. The first step to annotating TUs is to intersect the TU coordinates with the primary transcript coordinates. We separately considered TUs that overlap with single primary transcripts and TUs that overlap with multiple primary transcripts. For this analysis, we considered the identified primary transcript coordinates to represent a ‘ground truth’ annotation, and we only made minimal modifications to the primary transcript coordinates if they had very close overlaps, as defined by the user, with identified TUs. Regions of identified TUs that did not very closely match identified transcripts were assigned generic TU identifiers. The analyses described below are implemented by the single.overlaps() and multi.overlaps() functions, which are called by get.tu.gene.coords().

For TUs that overlap with single transcripts, we considered a reference case in which the transcript coordinates are contained within the boundaries of the TU (Figure S5a). We term this reference case ‘class 1’. This analysis evaluates whether the beginning of a TU is in close proximity to the beginning of the transcript. If this criteria was not met, the portion of the TU that is upstream of the gene annotation was assigned an arbitrary identifier and the identified TSS defines the start of a TU with named for the overlapping gene. Similarly, if the annotated transcript end was far from the boundary of the TU, then we set the end of the TU, named by the respective overlapping transcript, to the identified TTS. Here we would then annotate the downstream region of the TU with an arbitrary label. For example (Figure S5a), if the distance between the starts of transcript `X` and TU `X` is less than 200 bp, and if the distance between the respective ends is less than 1 kb, we set the identifier of TU `X` to transcript `X`. If the distance between starts is greater than 200 bp, we annotate the initial segment of the TU with a generic identifier (tu_class1_1, see main text Figure 1f and the associated browser session: https://genome.ucsc.edu/s/warrena%40virginia.edu/primaryTranscriptAnnotation_20190801). If the distance between ends is greater than 1 kb, we annotate the final segment of the TU with a generic identifier (tu_class1_2). Hence, we maintain a close correspondence between the identified transcript and an overlapping TU and we label marginal regions of transcription with generic identifiers.

Next we considered cases in which a TU was enclosed within a transcript (class 2, Figure S5b). Because this analysis was completed under the assumption that the identified TSS/TTS coordinates are veritable primary transcript boundaries, we simply extended the region of the enclosed TU to match the transcript. Then we implemented the same rules described for class 1 above. In this case, the algorithm will default to assigning the gene identifier to the TU, as the respective coordinates are identical.

We further considered cases in which TU-transcript overlaps were characterized by ‘overhangs’ (class3-4, Figure S5c,d). Again, here we extended the appropriate TU boundaries to match the corresponding transcripts such that no part of the transcript extended beyond the TU. We then applied the class 1 logic to assign TU identifiers based on the identified transcript coordinates.

For TUs that overlap with multiple transcripts, we considered a reference case in which all of the transcript coordinates were contained within the boundaries of the TU (class 5, Figure S6a). For the upstream-most and downstream-most overlaps, we applied analyses comparable to those for the class 1 scenario in which we used either the existing transcript annotation or introduce an arbitrary identifier depending on the proximities of the boundries to the TSSs and TTSs. If the regions between transcripts were sufficiently large, we introduced generic identifiers (e.g., tu_class5_1; see regions i1 and i2 in Figure S6a). As described above for single overlap classes 2-4, the logic implemented for class 5 could be applied to other scenarios of multiple overlaps (Figure S6b-d). For instance, in cases of upstream and/or downstream overhangs, the TU could be extended such that all overlapping transcripts were enclosed, then the class 5 logic could be applied to classes 6-8 (Figure S6b-d). See examples in main text Figure 1f.

## Supplementary results

### TSS and TTS identification

We evaluated the performance of `primaryTranscriptAnnotation` by comparing results obtained by using transcript coordinates inferred using our package against largest interval coordinates derived from GENCODE. The largest interval coordinates were obtained by taking the upstream-most TSS and the downstream-most TTS for each gene. To evaluate the performance of our TSS evaluation method, we examined the distances between read density peaks and inferred TSSs. For both annotations, the distributions of these distances showed a peak downstream of zero, corresponding to promoter proximal polymerase pausing (Figure S1a). However, we observed that the distribution for the inferred annotations showed a more focused peak with reduced variance (p = 2.1 × 10^*−*34^, Levene’s test) (Figure S1b,c). This shows that taking the longest of conventional mRNA annotations results in assigning transcription initiation sites to regions that are farther from pause sites than the corresponding distances based on our inferred TSSs.

We estimated TTSs using the novel method described above. We specified a distance beyond the most distal gene end annotation and evaluated whether there were intervening TSSs within this region. If TSSs were present, we clipped the region at the site of the most proximal TSS. Figure S2a shows the distribution of clip distances when we considered an interval of 100 kb beyond the most distal annotated gene ends. A prominent mode of clip distances is apparent at 100 kb, because many mammalian genes occur in tandem such that the end of one gene is close to the start of another. We used the clip distances to define a gene end peak search region for each transcript. We operationally defined TTSs as the points where spline fits to binned reads decayed to a specified fraction of the largest peak of the spline trace. Figure S3a,b shows examples of splines (blue or red) and estimated TTSs (vertical lines), in which the traces are shown throughout the TTS search regions. For example, Figure S3b shows the identified primary transcript coordinates for *Lypla1* and *Tcea1* (plus strand, red). The TTS for *Lypla1* was identified at the end of the search region because *Lypla1* immediately preceded the TSS for *Tcea1*. Note that the inferred TTSs typically extend beyond the largest interval TTSs (Figure S4d).

To characterize the differences between inferred primary transcript coordinates and largest interval coordinates, we evaluated the differences between the TSSs and TTSs of the two annotation sets. We found that 49% of the cases in which the TSS did not change, there was only a single exon 1, in which case our analysis was not designed to re-annotate the TSS. We found that 57% of the expressed genes with multiple exon 1 isoforms were re-annotated from the largest interval TSS to an inferred TSS. Figure S4a,b show the distribution of TSS differences. For the TSSs that were re-annotated relative to the largest interval, we found that 75% of the inferred TSSs are within 813 bp of the largest interval TSS (Figure S4b). As expected, the chance of being re-annotated increases as more exon 1 isoforms are characterized per gene (Figure S4c) As gene annotations progressively incorporate newer and more comprehensive TSS inference technologies, including 5′ RACE and CAGE-based approaches (Cvetesic et al., 2018; Leenen et al., 2016), our methods will facilitate context-specific TSS inferences with enhanced precision.

The distribution of TTS differences is shown in Figure S4d. We found that 75% of TTS differences are within approximately 15 kb, and in the majority of cases, the inferred TTSs are downstream of the largest interval TTSs (Figure S4e). These data show that inferred annotations result in substantially longer transcripts due to the inference of more downstream TTSs.

### Evaluation of transcript expression dynamics and RNA polymerase pausing

To determine whether applying inferred primary transcript coordinates leads to genome-wide variations in expression dynamics, we mapped the adipogenesis time-series PRO-seq reads to the inferred coordinates and projected the data onto principal components (PCs, Figure S7a). For comparison, we mapped the same data onto the conventional largest interval annotations and visualized the PC projections (Figure S7b). The results of this analysis show that the PC projections are nearly identical for both annotation sets. Thus, applying inferred primary transcript coordinates does not lead to genome-wide variations in expression dynamics, as defined by data projections onto PCs 1-3, which capture 83-89% of the variation in the data.

Next we addressed whether applying inferred primary transcript coordinates alters the results of differential expression analyses, as compared to the results obtained using largest interval annotations. To evaluate differential expression, we used a likelihood ratio test to determine whether transcript expression varies with respect to time over four hours. This analysis is analogous to using a 1-factor ANOVA to determine the effect of the time factor. We implemented this test for both inferred coordinates and largest interval coordinates. Figure S8a,b shows that the negative log values of the resulting false discovery rates (FDRs) tend to be higher for the expression levels quantified using inferred coordinates (i.e., lower FDRs; higher density above the unity line in Figure S8b). At a threshold of FDR < 0.001, we observed 11,351 genes that were differentially expressed for both annotations, 520 genes that were differentially expressed only for the inferred coordinates, and 241 genes that were differentially expressed only for the largest interval coordinates (Figure S8c). To test whether the counts of exclusive differentially expressed genes were identical for both annotations, we applied a binomial test of the null hypothesis that the proportions of the 761 exclusive genes (520+241) were identical (i.e., 50% each). The analysis was not consistent with the null hypothesis in which 50% of genes are proposed to be differentially expressed for each annotation (p < 2.2 × 10^*−*16^). This suggests that there are more differentially expressed transcripts after mapping read counts to inferred coordinates as compared to largest interval coordinates.

To determine whether the finding of elevated differential expression associated with inferred annotations was robust to the FDR threshold for defining significance, we varied this threshold and evaluated differential expression for both inferred and largest interval annotations. The number of differentially expressed genes observed for both annotations increased as the FDR was relaxed to larger values (Figure S8d, left). In contrast, the counts for exclusive differentially expressed genes decreased with respect to the FDR threshold (Figure S8d, right). Importantly, the counts of genes differentially expressed only for the inferred annotations were consistently higher than the corresponding counts associated with using the largest interval annotations. To evaluate the significance of this trend, we compared the results for the two gene annotations at each FDR using the binomial test as described above. We then applied the Benjamini-Hochberg correction to adjust the resulting set of binomial test p-values (Benjamini and Hochberg, 1995). The resulting set of adjusted p-values ranged from *p*_*adj*_ = 1.0 × 10^*−*35^ for a likelihood ratio test FDR threshold of 1 × 10^*−*6^ to *p*_*adj*_ = 4.6 × 10^−6^ for an FDR threshold of 1 ×10^*−*1^. These analyses demonstrate that mapping reads to inferred primary transcript coordinates results in improved sensitivity for the detection of differential transcript expression, as compared to the results observed when using conventional mRNA annotations.

Finally, we evaluated whether data-driven inferred co-ordinates enhance the sensitivity for detecting promoter-proximal polymerase pausing. For this analysis we considered expressed genes with inferred annotations, and we evaluated pausing across all pre-adipogenic time points. The inferred annotations were associated with elevated pause region composite profiles (Figure S9a). To examine the average precision of the paused regions associated with inferred versus largest interval coordinates, we plotted 0-1 scaled composites (Figure S9b). The results show that inferred annotations are associated with sharper pause region read distributions, on average, as compared to profiles based on largest interval coordinates. We computed pause indices for inferred and largest interval annotations (see Methods above). The results revealed that the distributions were significantly different based on applying the Wilcox test at each time point (FDR < 5 × 10^*−*140^, Figure S9c). These analyses demonstrate that the application of inferred coordinates enhances the precision for quantifying the position and degree of promoter proximal RNA polymerase pausing from genome-wide run-on data.

---

**Fig. S1.**
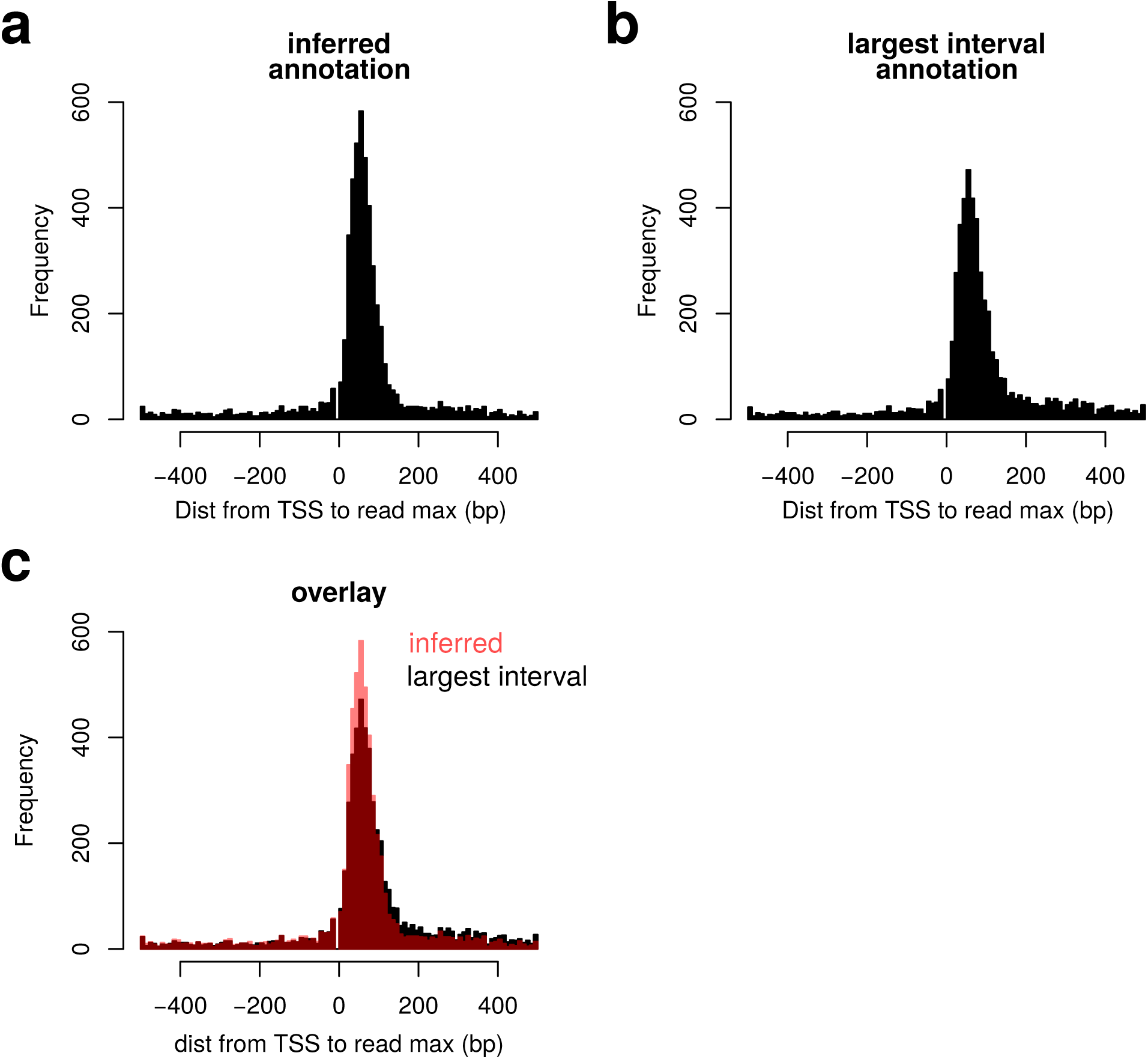
Inferred TSSs are more accurate, based on the expected distribution of paused RNA polymerase peaks. The distribution of distances between inferred (a) and largest interval (b) TSSs and regions of peak read density using a 1 bp sliding 50 bp window. (c) An overlay of the data from a and b indicate a more focused distribution of paused polymerase downstream of the TSS for the inferred start sites.

**Fig. S2.**
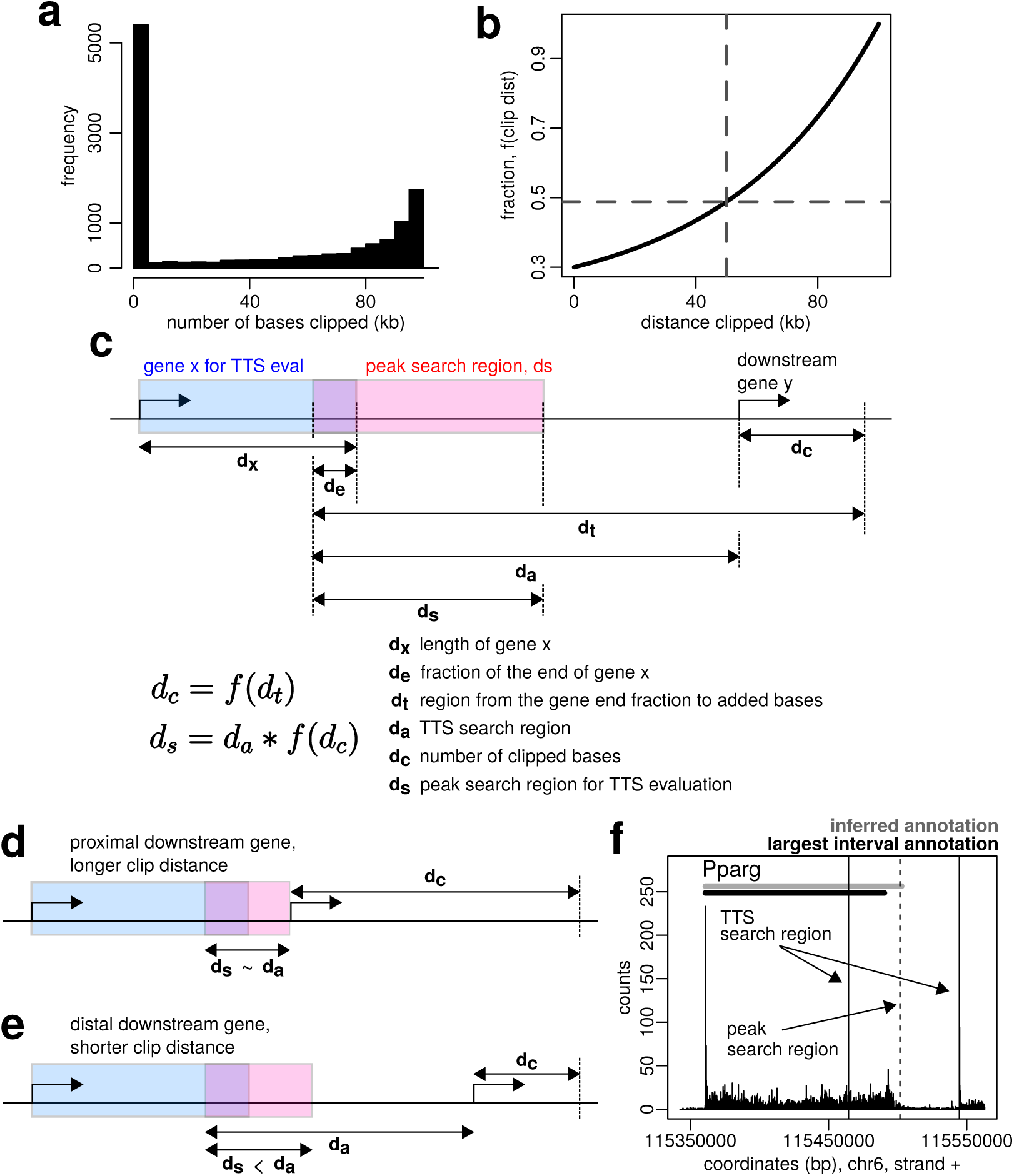
The TTS inference approach identifies transcriptional termination in a data-driven manner. (a) The distribution of clip distances is bimodal. The majority of genes are >100 kb from the the closest downstream gene (clip distance = 0). A subset of genes immediately tandem to adjacent genes; in such cases, ∼ 100 kb are clipped to avoid overlapping annotations. (b) The initial search region (*d*_*a*_) for the TTS is a function of the clip distance. (c) The terminology that is used to define the TTS search region and 3′ RNA polymerase peak search region is indicated in the schematic. (d) This example illustrates a scenario in which there is a TSS proximal to the end of the query gene on the same strand, so there is a large clip distance and a small peak search region. (e) Alternatively, a distal TSS results in a small clip distance and a relatively large peak search region. (f) Vertical lines indicate the TTS search region and the peak search for *Pparg*, along with the inferred annotation (gray bar) and the largest interval annotation (black bar).

**Fig. S3.**
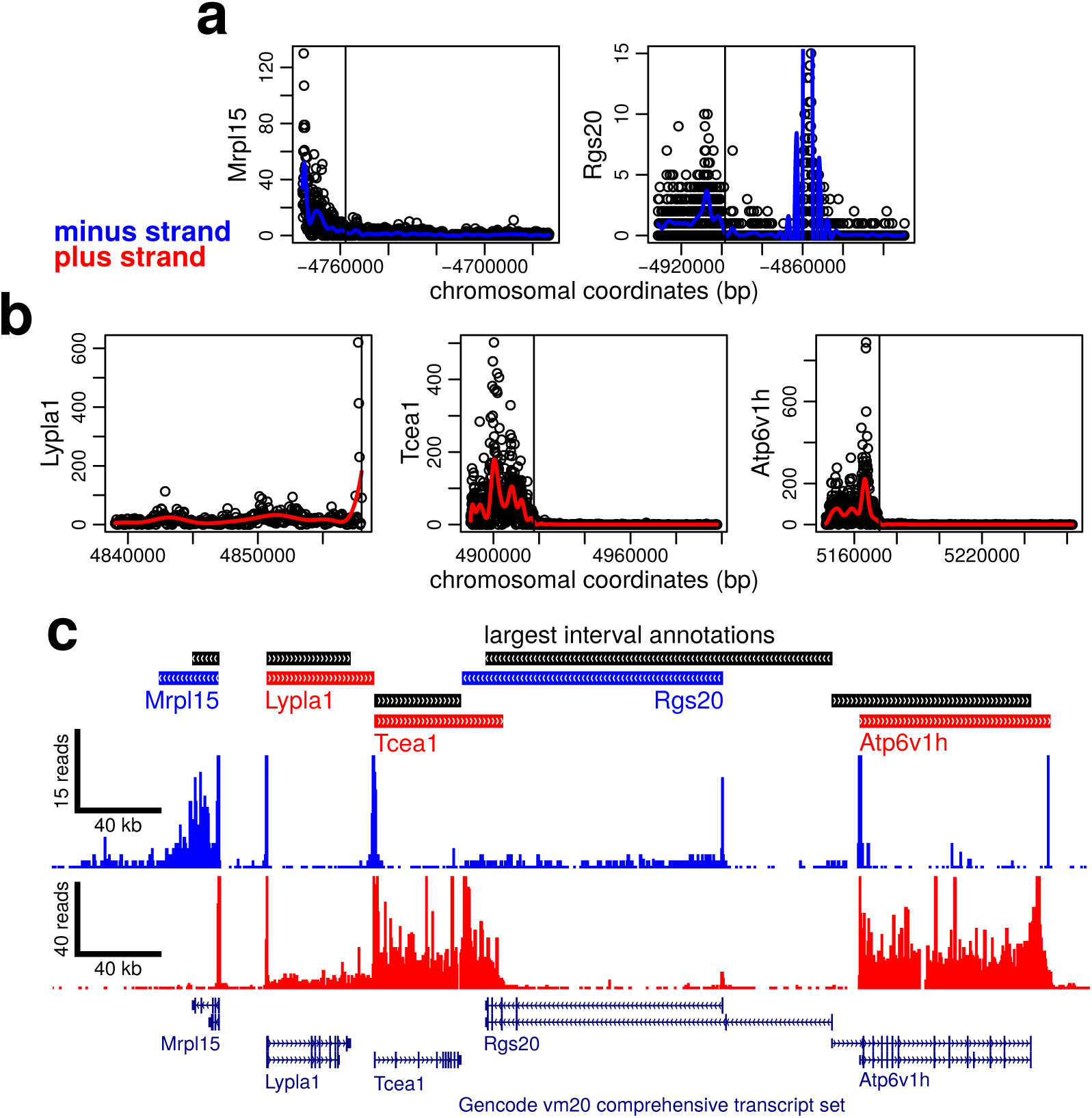
TTS inferences accurately reflect regions of apparent transcriptional termination. (a,b) Reads in the TTS search region were binned and quantified (black open circles) and smooth spline curves were fit (smooth curves). The black vertical lines indicate inferred TTSs. These vertical lines for Mrpl15, Rgs20, Tcea1, and Atp6v1h represent the point at which spline curves decay from an initial peak toward zero. The inferred TTS for Lypla1 was the end of the search region because Tcea1 is closely downstream and transcribed from the same strand. Negative numbers are used for the horizontal axes to indicate transcription from the minus strand. (c) Genome browser displays highlight the improved accuracy of the primary transcript annotations.

**Fig. S4.**
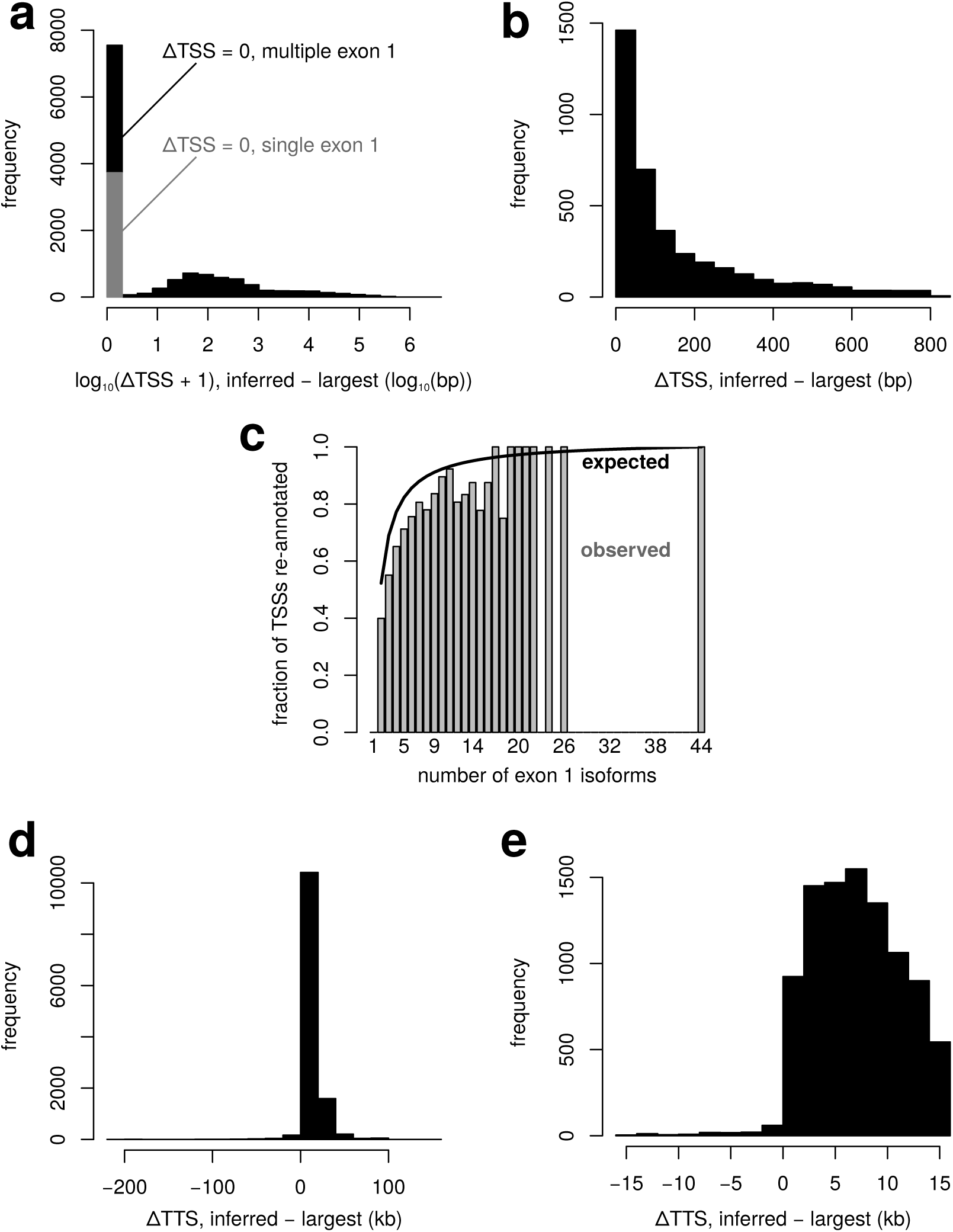
Inferred annotations produce longer transcripts due to distal TTSs. (a) The distribution of TSS differences between inferred and largest interval annotations. (b) The distribution of TSS differences up to the 75% quantile for those TSSs that were re-annotated relative to the largest interval. (c) A TSS is more likely to be re-annotated if more exon 1 isoforms are present in annotation files. (d) The distribution of TTS differences between inferred and largest interval annotations. (e) The distribution of TTS differences up to the 75% quantile.

**Fig. S5.**
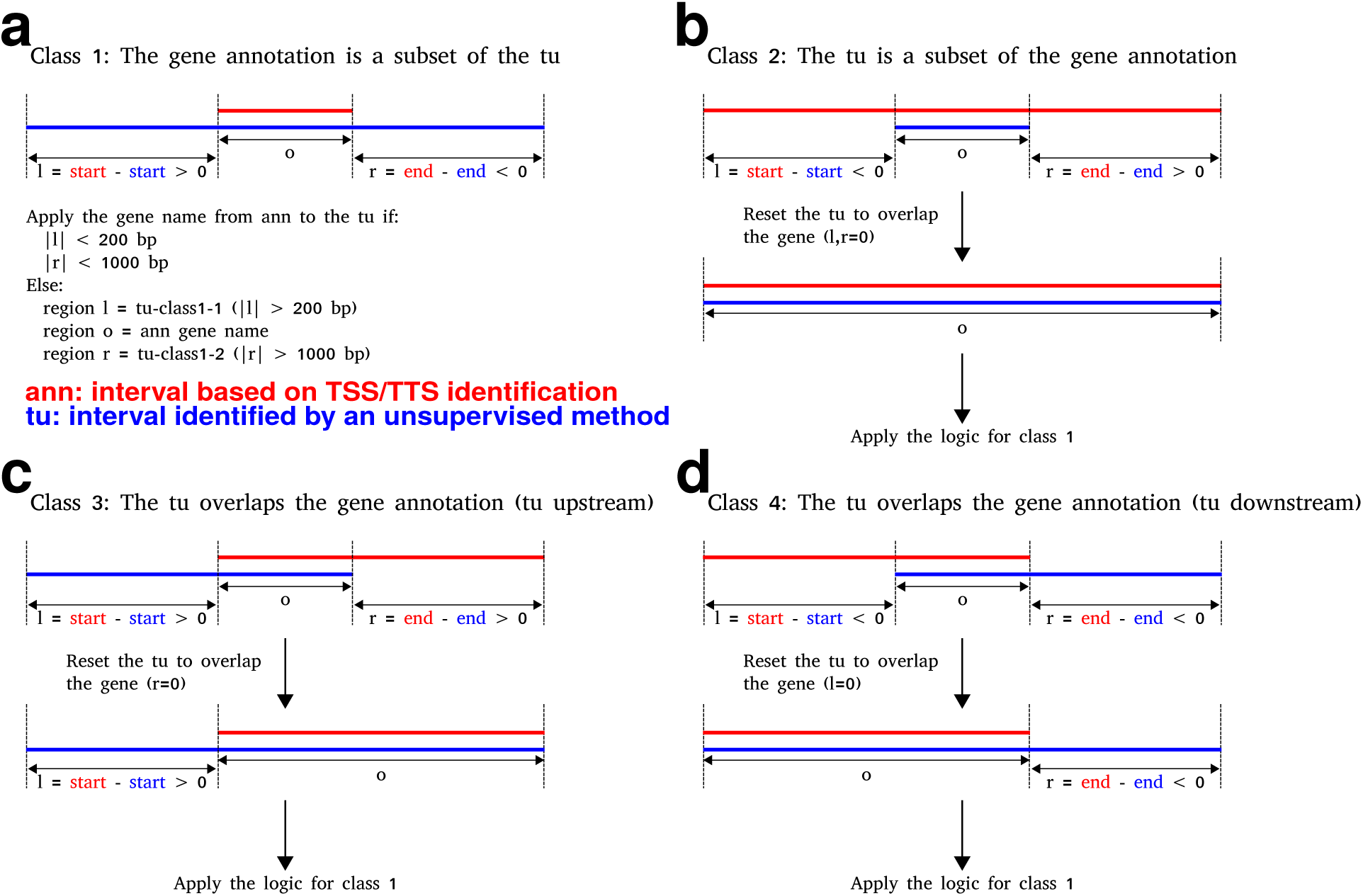
Rules for annotating TUs that overlap with a single transcripts. (a-d) The data-driven TSS/TTS intervals are used to redefine TU ends that fall within class 1-4 overlap profiles. Note that gene is shorthand for primary transcript.

**Fig. S6.**
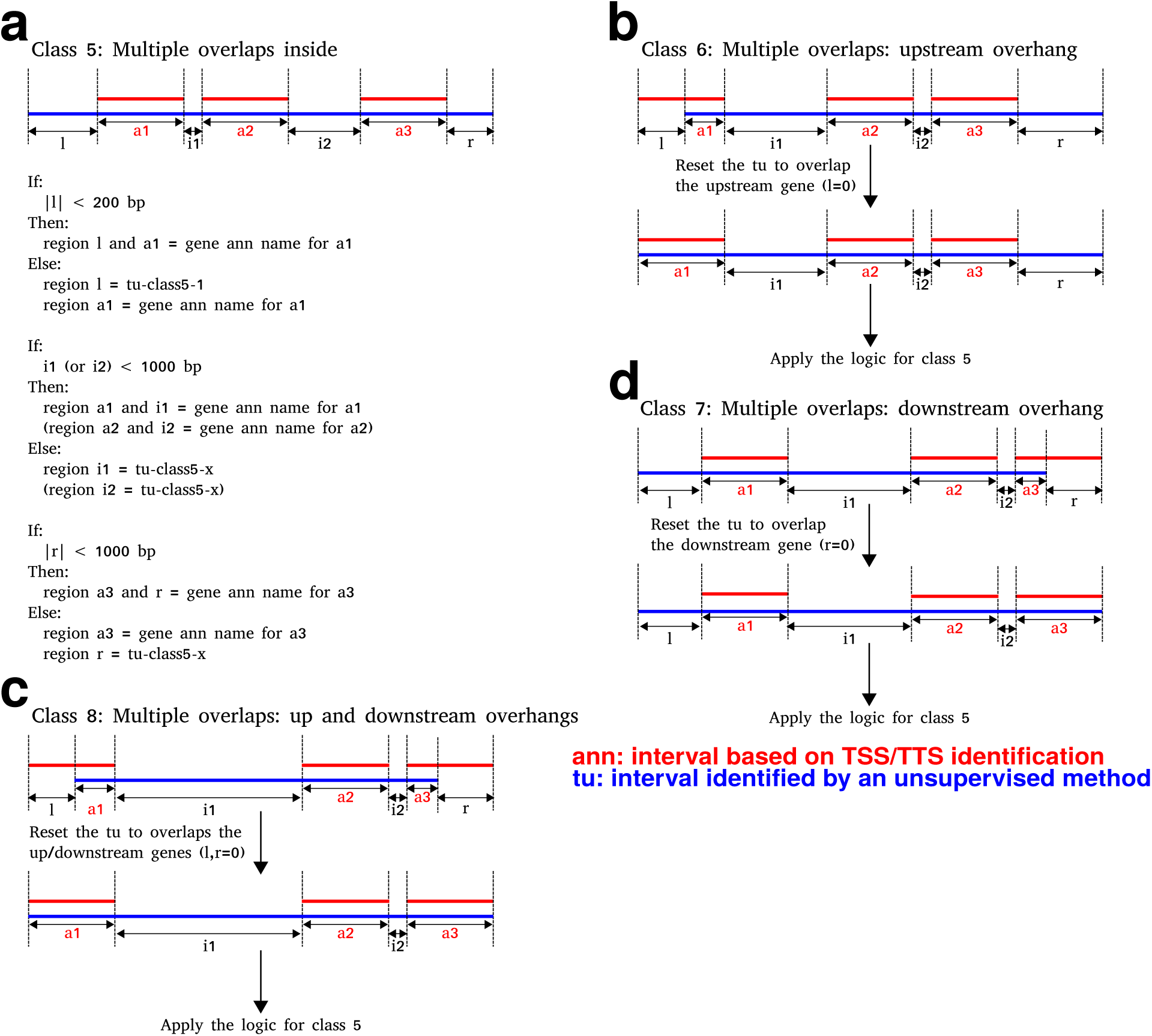
Rules for annotating TUs that overlap with multiple transcripts. (a-d) The data-driven TSS/TTS intervals are used to redefine TU ends that fall within class 5-8 overlap profiles. Note that gene is shorthand for primary transcript.

**Fig. S7.**
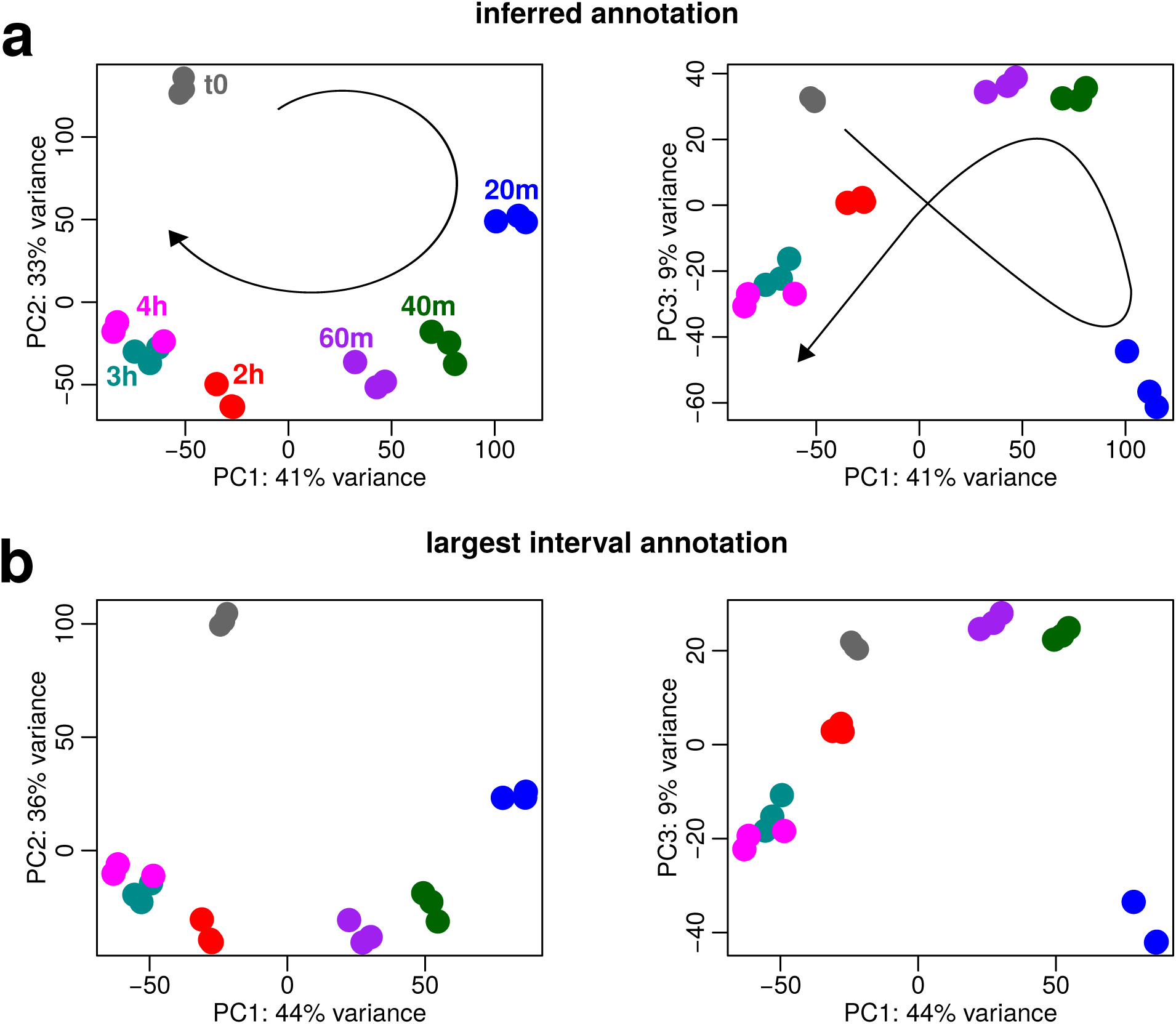
Principal component analysis of adipogenesis for inferred and conventional transcript coordinates. Principal component (PC) one and two (left), and PC one and three (right), projections for inferred transcript coordinates (a) are indistinguishable from largest interval coordinates PCA (b). The lines with arrows indicate the temporal trajectories of expression changes within the spaces defined by the PCs.

**Fig. S8.**
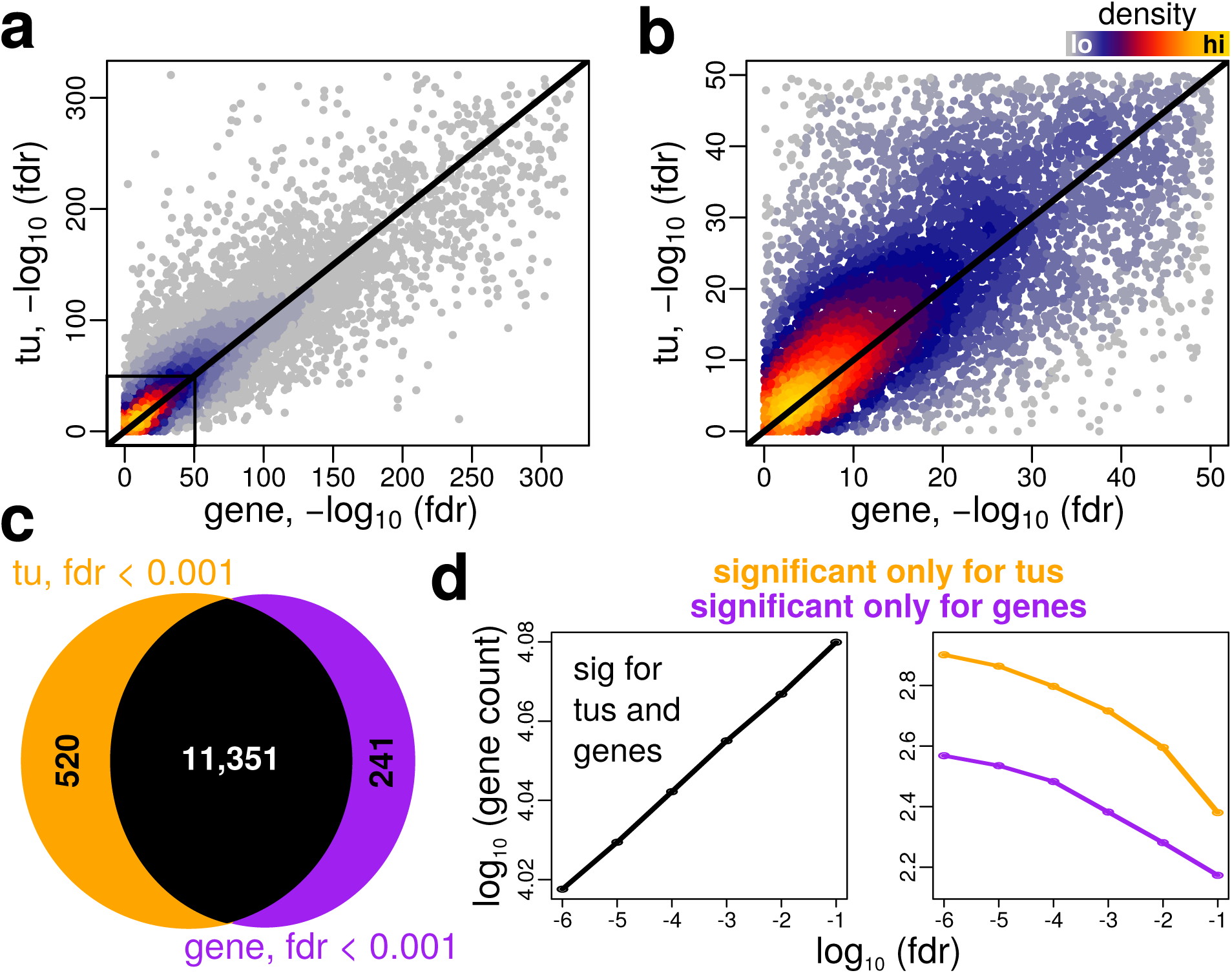
Inferred coordinates confer enhanced sensitivity of detecting differential transcript expression. (a) FDRs are correlated for inferred annotations (tu) and largest interval annotations (gene). (b) An expanded version of the left corner from panel (a) shows that FDRs tend to be more significant for inferred annotations (above the identity line). (c) At an FDR threshold of 0.001, there are 520 differentially expressed transcripts that are unique to inferred annotations and 241 that are unique to the largest interval annotations. Note that the Venn diagram is not drawn to scale for illustrative purposes. (d) TU annotations result in more significantly differentially expressed transcript counts over a range of FDR thresholds.

**Fig. S9.**
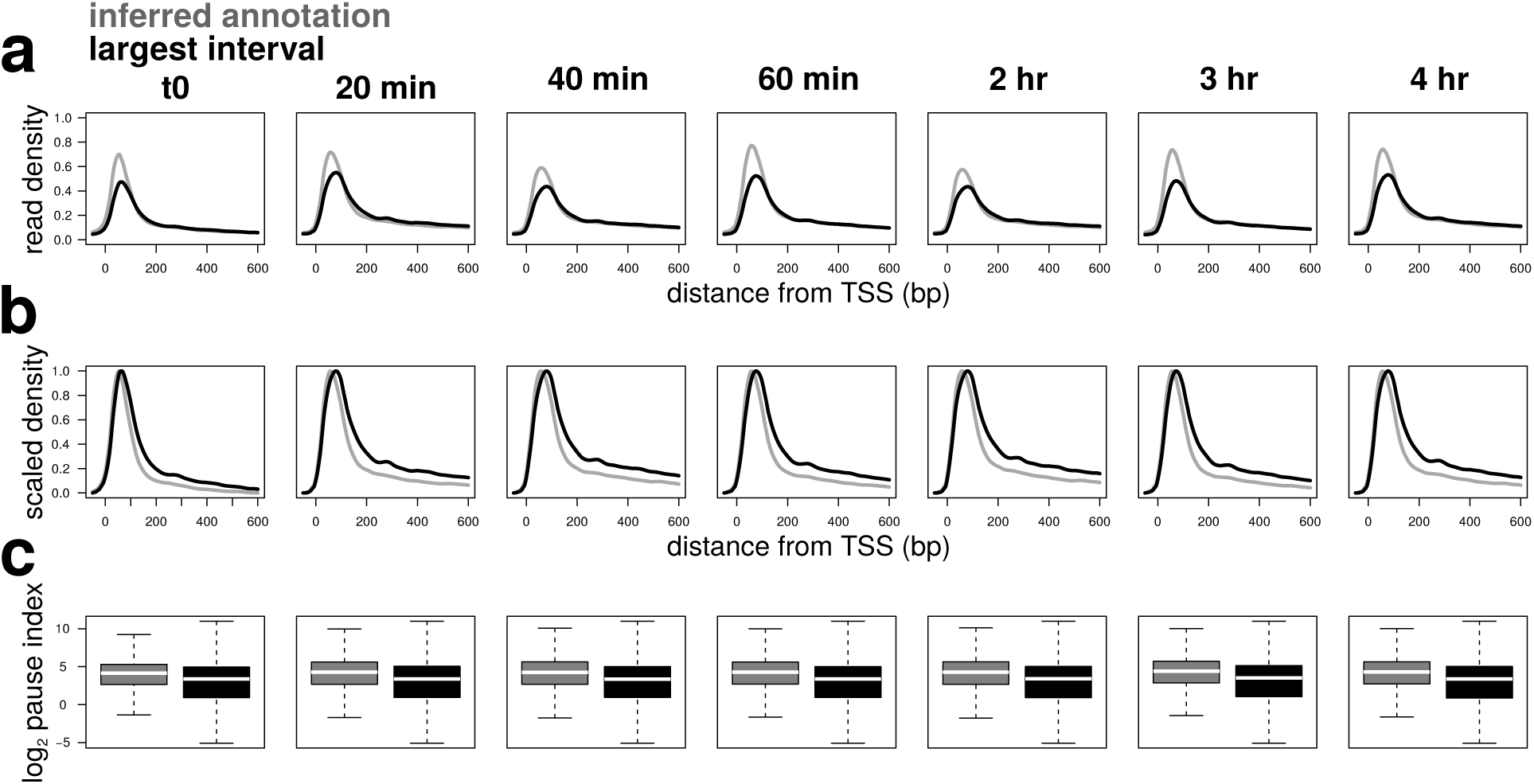
Inferred coordinates confer enhanced sensitivity for detecting polymerase pausing. (a) Pause region composite profiles are shown for both inferred coordinates and largest interval coordinates across pre-adipogenic time points. (b) Pause region composites are scaled to the interval 0,1 to emphasize the ‘sharpening’ observed for inferred coordinates. (c) Pause index distributions (TSS region / gene body read density) show elevated pause indices for inferred coordinates.

